# Towards Quantitative Imaging Biomarkers of Tumor Dissemination: a Multi-scale Parametric Modeling of Multiple Myeloma

**DOI:** 10.1101/613869

**Authors:** Marie Piraud, Markus Wennmann, Laurent Kintzelé, Jens Hillengass, Ulrich Keller, Georg Langs, Marc-André Weber, Björn H. Menze

## Abstract

The advent of medical imaging and automatic image analysis is bringing the full quantitative assessment of lesions and tumor burden at every clinical examination within reach. This opens avenues for the development and testing of functional disease models, as well as their use in the clinical practice for personalized medicine. In this paper, we introduce a Bayesian statistical framework, based on mixed-effects models, to quantitatively test and learn functional disease models at different scales, on population longitudinal data. We also derive an effective mathematical model for the crossover between initially detected lesions and tumor dissemination, based on the Iwata-Kawasaki-Shigesada model. We finally propose to leverage this descriptive disease progression model into model-aware biomarkers for personalized risk-assessment, taking all available examinations and relevant covariates into account. As a use case, we study Multiple Myeloma, a disseminated plasma cell cancer, in which proper diagnostics is essential, to differentiate frequent precursor state without end-organ damage from the rapidly developing disease requiring therapy. After learning the best biological models for local lesion growth and global tumor burden evolution on clinical data, and computing corresponding population priors, we use individual model parameters as biomarkers, and can study them systematically for correlation with external covariates, such as sex or location of the lesion. On our cohort of 63 patients with smoldering Multiple Myeloma, we show that they perform substantially better than other radiological criteria, to predict progression into symptomatic Multiple Myeloma. Our study paves the way for modeling disease progression patterns for Multiple Myeloma, but also for other metastatic and disseminated tumor growth processes, and for analyzing large longitudinal image data sets acquired in oncological imaging. It shows the unprecedented potential of model-based biomarkers for better and more personalized treatment decisions and deserves being validated on larger cohorts to establish its role in clinical decision making.

## 1. Introduction

Medical imaging of tumorous lesions is a means of choice for staging and monitoring patients with cancer. It enables early detection through population screening, permits to evaluate the growth of precursor lesions and assess qualitative and quantitative changes after clinical intervention (Fass, 2008). Although most imaging modalities allow for a volumetric quantification of lesions, current guidelines, like the Response Evaluation Criteria in Solid Tumors (RECIST) (Eisenhauer et al., 2009), are based on the manual assessment of the diametric size of a few lesions, for the sake of time, even if a large field of view image scan would have the potential to offer more and better information. To alleviate this problem, many tools for the automatic detection and segmentation of lesions are now becoming available for example in Positron Emission Tomography (Xu et al., 2017; Bieth et al., 2018) but also in Magnetic Resonance Imaging (MRI) (Kamnitsas et al., 2016) and Computer Tomography (CT) (Christ et al., 2016), due in particular to the advent of deep learning techniques in the medical imaging realm. They are bringing the full quantitive assessment of individual tumorous lesions and whole tumor burden, at each medical examination, within reach. This raises the question of how to properly analyze these data, both focusing on static and dynamical properties, a question of both scientific and clinical interest, which has recently started to be addressed (Hartung et al., 2017; Claret et al., 2018)

Numerous theoretical models of cancer evolution have been developed, grasping some of the complexity of the biological processes at stake, both for general aspects or more specific to a particular pathology. Descriptive models have in particular been derived for single lesions (Simeoni et al., 2004; Ayati et al., 2010; Herman et al., 2011; Gerlee, 2013; Benzekry et al., 2014; Murphy et al., 2016) as well as for the distribution of disseminated tumors (Iwata et al., 2000; Baratchart et al., 2015). But those are in general not well tested, due to the lack of observations at the right scale, or to the rarity of fully quantitative population datasets. Furthermore, the models describe different modes of disease propagation, and the crossover between different scales and regimes has been very little studied. In this paper, we establish a novel multi-scale approach, fusing local and global tumor growth models, at the onset of disease dissemination. We show how one can deal with different local growth patterns of lesions and analyze their dissemination in order to model the tumor load. We embed those descriptive models into a probabilistic framework, to deal with inference from observed data, and use that framework to compare different model options and therefore learn the local and global tumor growth models. We will finally show that this mixed analytical and statistical modeling approach can be used to extract model-based biomarkers, e.g. for patient stratification, that could ultimately serve as an objective tool for clinical evaluation.

We consider Multiple Myeloma (MM) as a case study, see box ‘Driving clinical problem’. MM displays a rapid development once the disease is manifest, but in the precursor phases of the disease, without myeloma related organ or tissue impairment, tumorous lesions and the tumor load are monitored via whole-body imaging over long time spans (Dimopoulos et al., 2015), alongside with serological and histological factors. It is therefore a typical example where proper image-based diagnosis and risk-assessment in the precursor states are crucial for making treatment decision (Ghobrial and Landgren, 2014; Ahn et al., 2015; van de Donk et al., 2016). This is posing hard problems in terms of analysis, and makes it a reference problem for empirical tumor growth modeling in disseminated diseases (Ghobrial, 2012). Our study will permit to validate key assumptions for the biological models at the organ and tissue level (Ayati et al., 2010) and for the dissemination process (Iwata et al., 2000), for the first time in a human population study.

## 2. Methods

Data-based tumor growth and disease progression modeling can be difficult, because available data are typically very limited in time. In particular, understanding free lesion growth, a cornerstone, is impeded by the occurrence of therapy, which usually begins soon after diagnosis. Animal models can help in that respect (Mehrara and Forssell-aronsson, 2014; Baratchart et al., 2015). But even with rather long time series, distinguishing between different growth models by individual curve fitting is very delicate (Murphy et al., 2016). Here we rely on microscopic biological models and translate them into clinically relevant observables: local lesion volumes and the tumor load. We propose a method to analyze longitudinal image time series of a population, in a hierarchical Bayesian framework (Ribba et al., 2012), both for lesion growth and tumor dissemination process. We also introduce covariates in a mixed-effect model (Lavielle, 2015), enabling statistical relevance tests of possible influential factors on the growth process.

### 2.1 Overview of the modeling approach

We analyze longitudinal data of a cohort of patients, as illustrated in Fig 1. At each observation time point, relevant features of all detectable lesions have been extracted, and used to compute the patient’s tumor load, as detailed in Sec. 3.1. Local lesions were also re-identified in follow-up scans, in order to gather a database of lesion growth time series.

**Figure 1:**
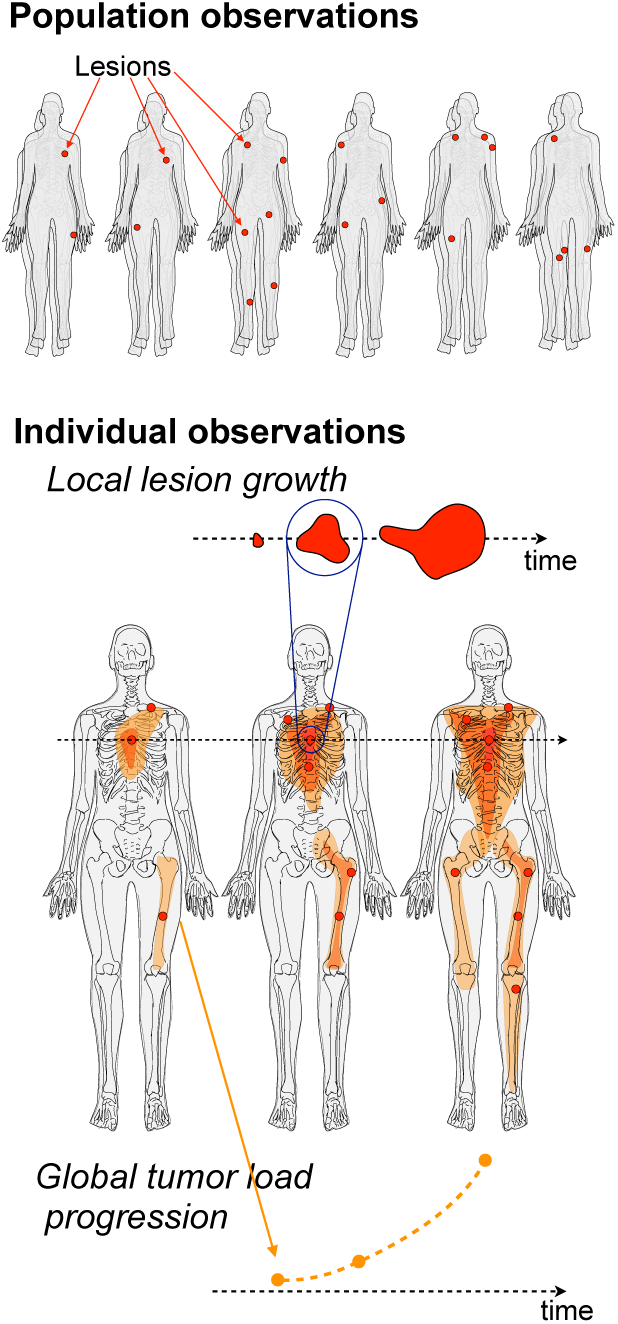
Overview of the approach: Observations and dataset.

##### Driving clinical problem

Multiple Myeloma is a cancer of plasma cells that is still incurable, with a median survival of 6 years at the time of diagnosis (Raab et al., 2009; Roellig et al., 2015; Kumar et al., 2017). It is a systemic cancer, which can be considered as a model for the cancer dissemination process (Ghobrial, 2012; Ghobrial and Landgren, 2014). Its development starts with the development and infiltration of clonal plasma cells within and into the bone marrow, homing into a niche and creating a micrometastasis. This initial cluster of malignant cells, can grow into a focal lesion and emit malignant cells that can in turn colonize other niches in the bone marrow. In overt MM, the malignancy causes endorgan damage, such as lytic bone lesions due to the perturbation of the bone remodeling cycle (Ayati et al., 2010).

The International Myeloma Working Group (IMWG) distinguishes two precursor stages, Monoclonal Gammopathy of Undertermined Significance (MGUS) and Smoldering Multiple Myeloma (SMM), preceding symptomatic Multiple Myeloma. This advanced stage is defined by the occurrence of end organ damage, following the CRAB criteria: ‘C’ for calcium elevation, ‘R’ for renal insufficiency, ‘A’ for anemia and ‘B’ for bone damage, corresponding to the appearance of bone lytic lesions on skeletal radiography or CT (International Myeloma Working Group, 2003). In 2014, further malignancy criteria were added to the definition of symptomatic MM, such as the presence of more than one focal lesions in MRI (Rajkumar et al., 2014; Rajkumar, 2016).

The incidence of MGUS is high in the population (1% of persons over 50 years-old (International Myeloma Working Group, 2003)) but only 15% of patients with MGUS will progress to MM (Kumar et al., 2017). Risk-stratification of patients in the early stages is therefore of primordial importance, to make treatment decisions (Ghobrial and Landgren, 2014; Ahn et al., 2015; van de Donk et al., 2016).

The functional-statistical modeling approach is schematically presented in Fig 2. In Sec. 2.2, we introduce our multi-scale mathematical modeling of disease progression. We first propose several biophysical models for the local lesion growth [*V*(*t*|*r, v*_0_), blue boxes in Fig 2], corresponding to different scenarii at the microscopic scale (Simeoni et al., 2004; Ayati et al., 2010; Gerlee, 2013; Benzekry et al., 2014; Murphy et al., 2016). We also propose an effective model for the crossover from the single-lesion regime to the dissemination regime [*V*_tot_(*t*|*R, V*_0_), green box in Fig 2], which builds on the local lesion growth model, as well as on the IKS model, a model for the dissemination process (Iwata et al., 2000).

**Figure 2:**
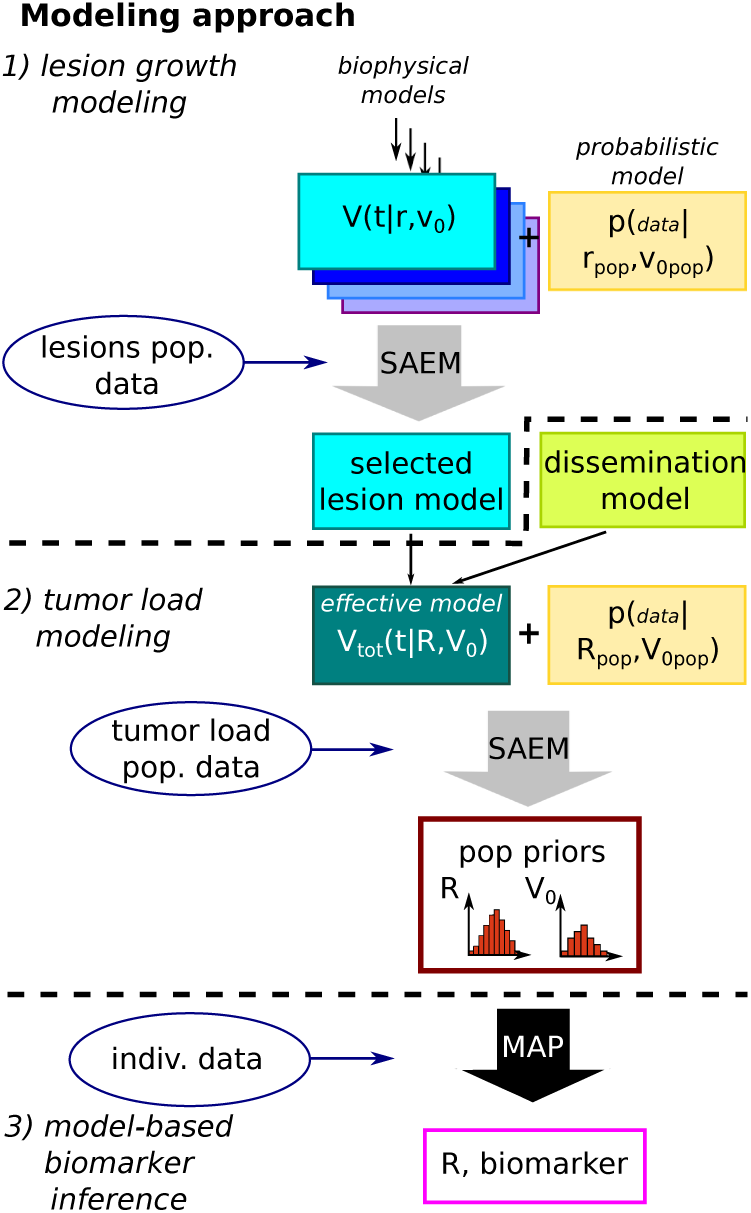
Overview of the modeling approach (SAEM stands for Stochastic Approximation of Expectation Maximization and MAP for Maximum a posteriori estimator).

Another necessary component to compare models with measurements, is the probabilistic model, which encodes how observations are generated from the predictive model [yellow boxes in Fig 2]. As detailed in Sec. 2.3, we rely on mixed-effects models (Lavielle, 2015), that are adapted to population-based tumor growth modeling (Bastogne et al., 2010; Ribba et al., 2012; Hartung et al., 2014; Baratchart et al., 2015)(Benzekry et al., 2016; Claret et al., 2018). We use a proportional error model, to account for a measurement error that increases with the size of the lesion. We also use population priors for the model parameters to increase the statistical confidence on shorter time series, and to incorporate covariates into the predictive model.

The confrontation of each mathematical model with the corresponding dataset, and the estimation of the unknown model parameters, is made via the Stochastic Approximation of Expectation Maximization (SAEM) algorithm (Delyon et al., 1999; Kuhn and Lavielle, 2004; Samson et al., 2007) [gray arrows in Fig 2] introduced in Sec. 2.4 and we use the implementation from the Monolix software (Monolix). It learns the population parameters from the data. To quantify the accuracy of the predictive model on the observed data, we compute the log-likelihood. Model comparison is then performed thanks to the Bayesian and Akaike Information Criteria (BIC and AIC) and assessed with bootstrapping. Covariates are further tested with the Wald and Likelihood ratio (LR) tests, see ‘Supplementary Method 1’. From the selected model and corresponding population priors, the individual parameters of a time series can be computed with a Maximum a posteriori estimator [black arrow in Fig 2], cf Sec. 2.4. We propose to use them as biomarkers, and for SMM patients, we explicitly use the tumor load growth rate. Its power for risk-stratification is assessed with Receiver Operating Characteristic curves (Zweig and Campbell, 1993), Kaplan-Meier survival plots (Kaplan and Meier, 1958) and log-rank tests (Peto et al., 1977), as presented in ‘Supplementary Method 2’.

### 2.2 Mathematical parametric models

The central elements of our modeling approach are descriptive functional models. We base our analysis on existing microscopic models of tumor and lesion growth in general (Simeoni et al., 2004; Gerlee, 2013; Benzekry et al., 2014; Murphy et al., 2016), as well as specific models for MM (Ayati et al., 2010; Herman et al., 2011), and for the dissemination process (Iwata et al., 2000). We aim at deriving tractable models of disseminative disease in a joint framework, and our approach is summarized in box ‘Descriptive tumor load model’.

#### 2.2.1 Local lesion growth models

The lowest scale of our modeling approach is given by individual tumors, also called focal lesions in MM. In this paper we assume that all lesions follow the same parametric growth model, i.e. that the involved biological processes are the same. This is reasonable as the lesions are all developing in the bone marrow. But different lesions potentially have different parameters, e.g. initial volume and growth rate, that may depend, for example, on the local environment of the lesion (Kumar et al., 2017) as well as its subclonal mutation status (Rasche et al., 2017). We introduce below several general tumor growth models, which are biologically founded (Simeoni et al., 2004; Gerlee, 2013; Benzekry et al., 2014; Murphy et al., 2016), as well as a more specific one for the microscopic biology of MM focal lesions (Ayati et al., 2010), which we interpret at the macroscopic scale of interest. All models are later confronted with observations.

##### Linear growth

We first introduce the linear growth model, in which the volume of a lesion can be written as:

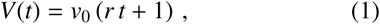

where *v*_0_ is the initial volume and *r* the growth rate, e.g. in month^−1^. This simple model corresponds to a constant rate growth, and holds at the later stage of tumor growth in some cases (Simeoni et al., 2004; Benzekry et al., 2014; Murphy et al., 2016).

##### Cubic growth

A cubic growth corresponds to a rate of change of the volume proportional to the surface area of the tumor itself, in a spherical approximation, 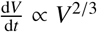,and reads

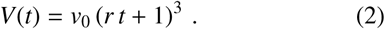

This assumes that only the surface of the tumor is actively participating in the growth, which is justified in the case of a solid tumor eroding its environment at its border (Herman et al., 2011; Gerlee, 2013; Murphy et al., 2016), representing a plausible model for lytic bone lesions.

##### Exponential growth

Another important tumor growth model is the exponential growth,

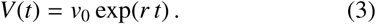

It corresponds to a volumic rate of change proportional to the volume, 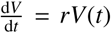. It therefore assumes that all cells of the tumor participate in its growth in the same manner, and typically holds in the early stages of tumor development (Gerlee, 2013; Benzekry et al., 2014; Murphy et al., 2016).

##### Diffusive growth

Ayati *et al.*, have derived in Ref. (Ayati et al., 2010) a mathematical model for bone remodeling and lesion growth in MM. Their modeling is based on a diffusion equation with a Gompertz-like saturation term for the local tumor density. In ‘Supplementary Method 3’, we show that it can be reduced to a two-parameters growth model for the tumor volume,

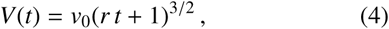

which corresponds to a half-cubic growth law, lying between the linear and cubic growth. Such a growth law can also be justified by advanced considerations on the fractal nature of the tumor vasculature (Herman et al., 2011; Benzekry et al., 2014).

Those models, Eqs. (1)-(4), comprise two free parameters each, *v*_0_ and *r*, which have a clear biophysical interpretation. There exists more involved models describing individual tumor growth with three parameters, for example the Gompertz and the van Bertalanffy models (Gerlee, 2013). Even more complex, are the models from Refs.(Herman et al., 2011) and (Ribba et al., 2012), which consider metabolic processes in details, but result in a high number of free parameters. For the sake of simplicity, and to avoid overfitting, given the limited length of the available time series, we restricted ourselves to the above models with two free parameters.

#### 2.2.2 Global tumor load models

The tumor load is obtained by summing up the volumes of all tumors detected in one patient, *V*_tot_ = Σ_lesions_ *l*’ *V*_*l*’_. We now consider how the local lesion growth models introduced above translate to this global scale. Let us distinguish between the volume of the putative initial lesion, *V*_init_, which is the largest detected lesion at first observation time, and the volume of all subsequent disseminated lesions, *V*_diss_:

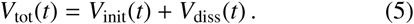

Note that we single out the first detectable lesion for modeling purposes, but this does not necessarily mean that it is the strict origin of the cancer. In the following, we simplify MM complex biology and assume that the cancer propagates from this leading cluster of cancerous cells, which grows and emits malignant cells, at a rate which depends on its size. Those in turn have a chance to settle in other locations, and give rise to further cell-emitting growing clusters, gradually making up the disseminated burden.

##### Early regime: initial lesion dominated

In the early stages of the disease, the initial cluster of malignant cells is the most prominent one and the disseminated burden *V*_diss_ can be neglected, such that

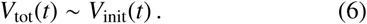

In that case, the tumor load growth model is the same as the lesion growth model in Eqs. (1)-(4).

##### Later regime: dissemination dominated

The Iwata-Kawasaki-Shigesada (IKS) mathematical model (Iwata et al., 2000) for metastasis formation seems particularly well suited to describe the evolution of MM. It assumes that growing tumors emit malignant cells at a rate proportional to the fraction of tumors cells in contact with blood vessels *w*(*v*) = *mv*^*α*^, where *v* is the volume of the tumor and *α* a fractal dimension related to the vascularization of the tumor. Similarly as above, *α* would be equal to 1 if the whole tumor volume can emit malignant cells, and *α* = 2/3 if only the surface of a round-shaped tumor can emit malignant cells. Emitted malignant cells can develop into new lesions far away from the original site, which will grow following the same tumor model, and emit further malignant cells. Note that the model assumes that each lesion grows with the same parameters, which is not strictly the case, as they depend on the lesion’s microenvironment and its subclonal mutation status (Kumar et al., 2017; Rasche et al., 2017). However, as extra-medullar dissemination is rare in SMM, their growth conditions should be more homogeneous than in multi-organ metastasis.

In ‘Supplementary Method 4’, we recall the solutions of the IKS model, as derived in Refs. (Iwata et al., 2000; Struckmeier, 2003; Evys et al., 2009), and show that for the different tumor growth models described above, the number of disseminated lesions *N*_diss_(*t*) and the disseminated burden *V*_diss_(*t*) asymptotically displays an exponential behaviour. Such that, as soon as the volume of the initial lesion becomes negligeable, we expect

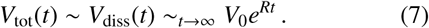

##### Crossover regime: effective modeling

In the crossover regime, neither of the terms of Eq. (5) is negligeable. In the cases where *V*_init_(*t*) follows a power-law, [Eqs. (1), (2) and (4)], *V*_tot_(*t*) will effectively appear as a power-law with a higher power than *V*_init_(*t*), due to the exponential behaviour of *V*_diss_(*t*). We therefore propose the effective model

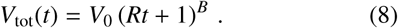

### 2.3 Probabilistic modeling: Mixed-effect model

To describe inter-subject variability in the population and compare parametric models with available longitudinal data, we need a probabilistic model. Here we use nonlinear mixed-effects models (Lavielle, 2015), which are adapted to population-based tumor growth modeling (Bastogne et al., 2010; Ribba et al., 2012; Hartung et al., 2014; Baratchart et al., 2015). In each experiment, we consider a dataset of observations {*y*_*i j*_} made at time *t*_*i j*_, where *j* is the index of the time series (lesion or patient in the following), and *i* is the index of the observation point. We are then comparing observations with the predictive growth model, *V*(*t*|*θ*), where *θ* = (*v*_0_, *r*) and (*V*_0_, *R*) for the lesion and the tumor load modeling, respectively. For that, using population priors permit to increase the statistical confidence on shorter time series, and to incorporate covariates into the predictive model.

#### Residual error model

The probabilistic model includes a discrepancy between the predictions and the observations, which can either stem from the inaccuracy or incompleteness of the parametric model as well as from the imperfections of the measurement process. To account for it, we assume that the measurements are normally distributed around the model predictions, following a proportional error model:

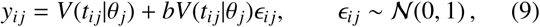

where *b* is the error parameter. We have tested other standard error models (constant, combined and power-law error models). They systematically gave worse results on all experiments (i.e. lower BIC values).

#### Population priors

To introduce variability, time series are considered independent from each other, and the model parameters *θ* are taken as log-normally distributed among them. Note that *θ* represents initial volumes and growth rates in this paper, which are indeed expected to take strictly positive values. We use

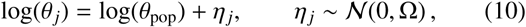

which is parametrized by *θ*_pop_, a vector of median values, and a diagonal matrix Ω = diag(ω_*θ*_), where ω_*θ*_ is also a vector, representing the correlation matrix of the logarithm of subject parameters *θ*.

#### Categorical covariates

Categorical covariates split the subjects into different groups, such as the sex of the patient or the region of the lesion location. To study their influence on the prediction, we include them in the prior, in a so-called mixed-effects model (Lavielle, 2015). In the case of a categorical variable that takes *K* values *k* = 0, 1, …*K* − 1, we write

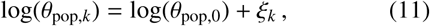

where *ξ*_0_ = 0 for the reference group *k* = 0.

### 2.4 Parameter inference: Stochastic Approximation of Expectation Maximization

We have introduced above a probabilistic model, which describes how data can be generated from the parametric model. We are left with the task of evaluating the best set of population parameters Ψ = (*θ*_pop,0_, {*ξ*}_*k*=1…*K*−1_, ω_*θ*_, *b*) and subject parameters 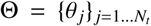, where *N*_*t*_ is the total number of time series, to match the observed dataset.

#### Population parameters

In the Maximum likelihood estimation (MLE), one searches the optimal population parameters as

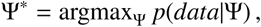

where *data* refers to all observations {*y*_*i j*_|*t*_*i j*_} that are available. One has 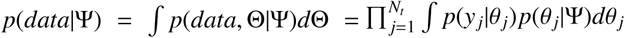, where *p*(*y*_*j*_|*θ* _*j*_) is determined by Eq. (9) and *p*(*θ* _*j*_ Ψ) is given by Eqs. (10)-(11). We use the SAEM algorithm to estimate Ψ^*^ (Delyon et al., 1999; Kuhn and Lavielle, 2004; Samson et al., 2007), which is a stochastic version of the Expectation Maximization algorithm, in which the computation of the expectation of the data in the ‘E step’ is replaced by a stochastic approximation. We use the implementation of the Monolix software and the nlmefitsa function in Matlab (Monolix; Matlab).

#### Log-Likelihood

To estimate the accuracy of the resulting regression over all time points of all time series, we compute the log-likelihood

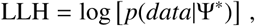

which is a quantification of the adequacy of the model on our data. Following Refs. (Kuhn and Lavielle, 2005; Samson et al., 2007) we use an importance sampling estimation, as implemented in Monolix and Matlab (Monolix; Matlab).

#### Individual parameters

Parameters for individual time series are estimated with the maximum a posteriori estimator. It consists in searching 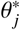, for each time series *j*, to maximize

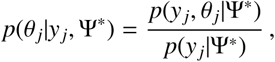

which is estimated using a Markov chain Monte Carlo (MCMC) method, also provided in Monolix and Matlab.

## 3. Experiments and results

In this section, we present the multi-scale modeling of disease progression in the SMM state. We compare several biologically-founded models for the growth of individual lesions, as well as for the tumor load, by confronting them with the datasets. We also carry out an analysis of the influence of the patient’s sex and lesion location on the growth rate. We finally propose to consider the extracted parameters as model-based biomarkers, and we assess their predictive power for the transition to MM.

### 3.1 Tumor imaging data

Our datasets result from the analysis of a large cohort of 63 SMM patients from the University Hospital of Heidelberg and the German Cancer Research Center, that both follow the same protocol for imaging and treatment decisions. We focus on the MRI modalities, which directly image focal lesions and the tumor load and is used to detect rapid progression to MM (Ghobrial and Landgren, 2014), whereas lytic bone lesions only appear at a later stage on CT scans. Our work follows a study for volumetry based biomarkers (Wennmann et al., 2018) in whole-body MRI scans. We use the same cohort and additionally analyzed all non whole-body scans that were available for those patients. In total, over 370 MR volumes were analyzed, with a median time interval of 1.1 years between scans, and a median patient follow-up time of 5.9 years. All detectable focal lesions were manually volumetrized, as illustrated on a T1-weighted sequence in Fig. 3, and tracked in time, as sketched in Fig 1, before the occurrence of the CRAB criteria or any systemic therapy. The manual volumetrization was performed by a research assistant with medical training, under the supervision of an experienced musculoskeletal radiology resident. When both T1- and T2-weighted sequences were available, the latter with fat suppression, the volume of the lesion was quantified in both modalities and averaged as in Ref. (Wennmann et al., 2018). This resulted in the detection of 180 lesions in 33 patients, each being observed at 2.19 different time points on average (8 time points for the longest observation).

**Figure 3:**
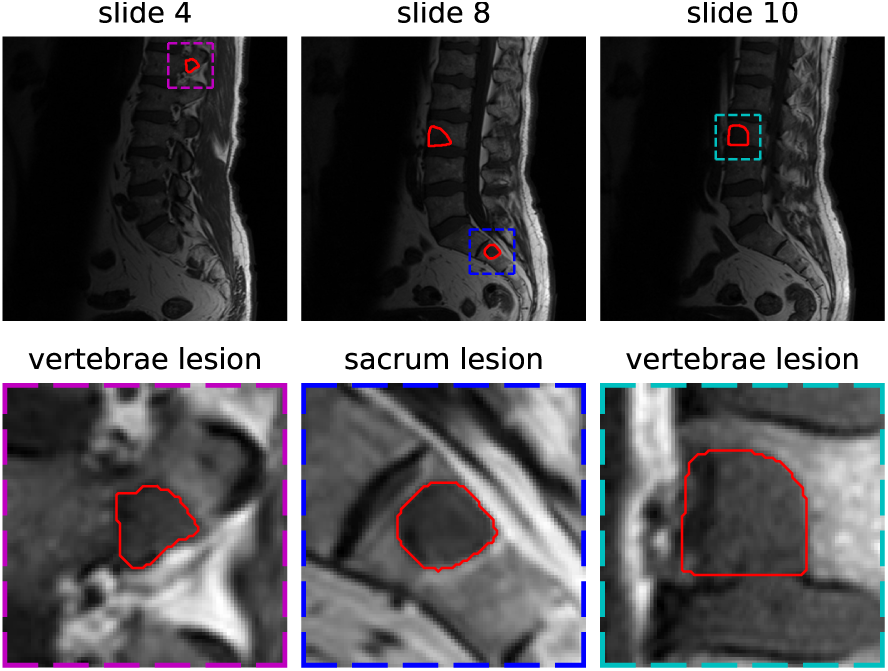
Three-dimensional segmentation and volumetrization of all visible hypo-intense focal lesions in a T1-weighted MRI sagittal MRI sequence.

##### Descriptive tumor load model

The tumor load is made of the putative initial lesion and the disseminated burden representing the process of new lesions starting to grow and to contribute to the overall load:

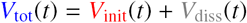

*V*_init_(*t*) ∈ {linear, cubic, diff., exp.} and *V*_diss_(*t*) ∼ exp.

We propose an effective model in the observation range:

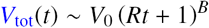

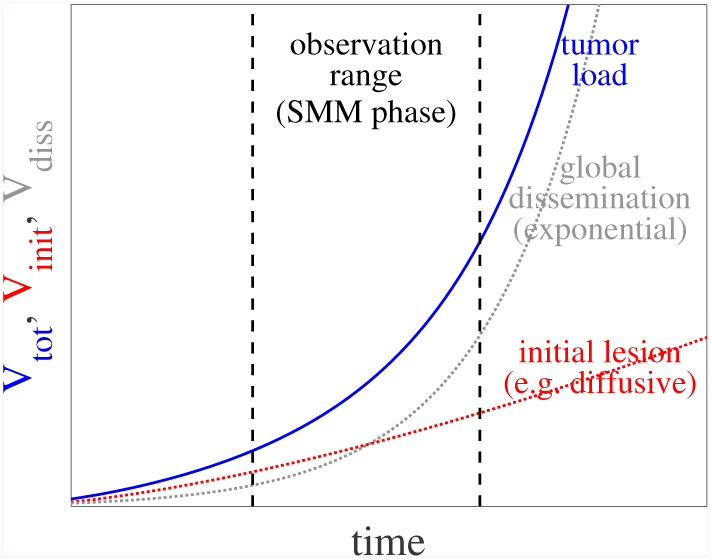

#### Focal lesions series

Among those focal lesions, 49 were detected at 3 or more different points in time. 36 of this subset originate from 11 male patients (M), and the 13 others from 8 female patients (F). The location of the tumor was also recorded and classified into 13 different anatomical regions (see ‘Supplementary Table 1’). Those 49 series constitute the first dataset, with 3.96 time points per series on average, which is used for lesion growth modeling.

#### Tumor load series

We also derived the total tumor load for each patient, by summing up the volume of all detected lesions at each time point. In rare cases, focal lesions became too diffuse to be properly segmented or were targeted by a local therapy, like radiotherapy. As the tumor load is not expected to decrease in the absence of therapy, if a tumor had already been detected but could not be segmented, we filled-in the data with its last measured volume. Selecting the series with 3 or more time points, we constitute a dataset of 21 patient series (13M and 8F) bearing 1 to 16 tumors (median of 4) with an average of 4.48 time points per series and a median total observation time of 3.7 years. The analysis of those tumor load series is used for tumor load modeling.

#### Progression to MM

For the 26 patients with focal lesion measurements on at least two different time points, we create a tumor load dataset, as above, that we complement with the date of transition to the progressive state of MM, as defined by the CRAB-criteria from the IMWG (International Myeloma Working Group, 2003), the standard procedure until 2014. If the transition did not occur, the data is censored with the date of the last information about the patient. We use this dataset for a risk-analysis of SMM patients with focal lesions, comparing different radiological biomarkers.

### 3.2 Modeling local lesion growth

#### Lesion growth model

Using the focal lesions dataset, we estimate the population parameters of a mixed-effect model without covariate, as presented in Sec. 2.1 with the different growth models introduced above, using SAEM. We compare a linear, a cubic, an exponential and a diffusive growth model [Eqs. (1)-(4)]. From the first column of Table 1, which presents the resulting BIC values for each model, we conclude that the diffusive growth model is selected, as it has the lowest BIC. It gives slightly better results than the linear and the cubic models (mean BIC separated by 5 resp. 13 confidence intervals), and much better than the exponential growth model. The learned population parameters for the diffusive model are *r*_pop_ = 3.2(0.9) × 10^−2^ month^−1^ and *v*_0,pop_ = 393(72) mm^3^, associated with the error parameter *b* = 0.24(0.02), where the number between brackets is the standard deviation estimated from the Fisher information matrix. The predictions from the diffusive model are shown in Fig 4 for two patients, and further lesion time series are presented in ‘Supplementary Figure 1’.

**Table 1:** Mean and standard deviation of BIC values for different lesion growth models, in two mixed-effects models: without covariate (first column) and with the patient’s sex as a covariate for *r* (second column).

| Model: | no covariate | M/F cov. for $r$ |
| --- | --- | --- |
| Linear | 2951.89 (0.12) | 2952.99 (0.13) |
| Cubic | 2954.65 (0.17) | 2956.99 (0.19) |
| Exponential | 2974.77 (0.22) | 2976.85 (0.21) |
| Diffusive | 2950.33 (0.17) | 2952.96 (0.17) |

**Figure 4:**
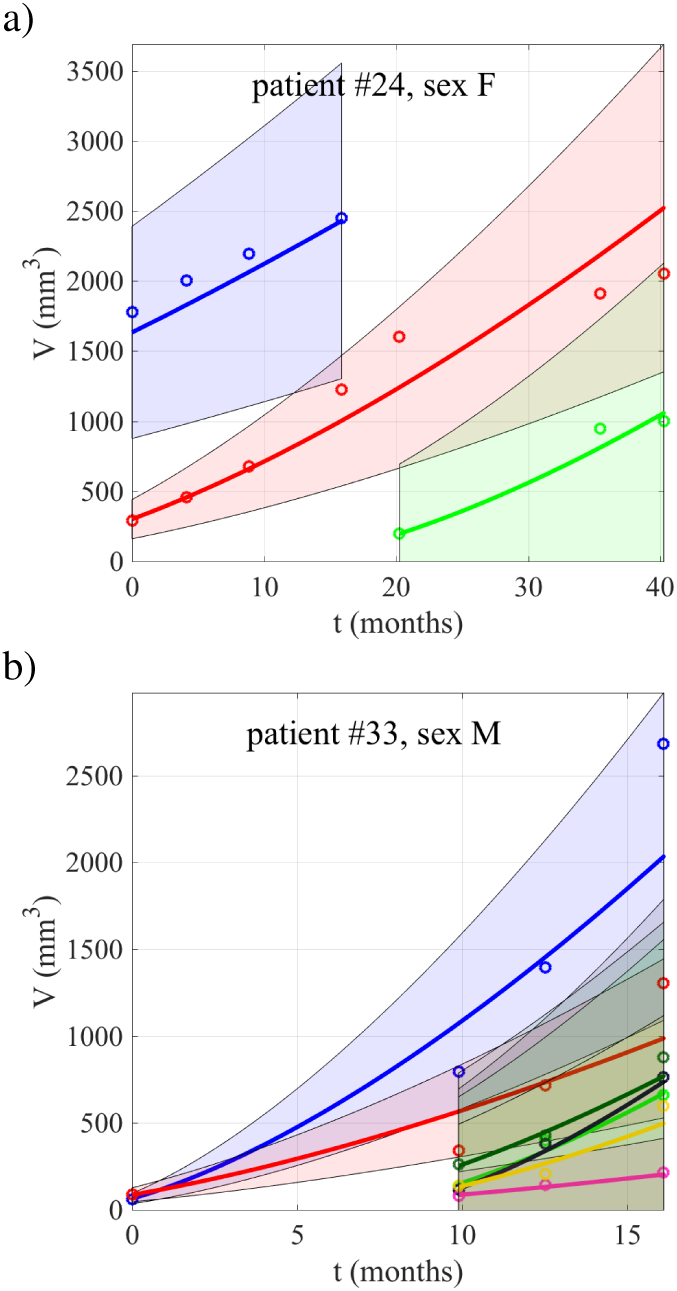
Local lesion growth time series (dots) with the predictions (solid lines) and the corresponding 95% error margin (shaded area), for a female and (b) a male patient. The predictions are based on the SAEM results using the diffusive parametric growth model, Eq. (4), and no covariate.

To assess the robustness and the generalizability of our model selection, we use a bootstrapping approach to resample our dataset, and compare the two best models selected above, whose BIC values are closer together than the other models. Due to the difficulty of resampling observations when doing regression (Davison and Hinkley, 2013), we use case bootstrapping to preserve inter-series variability (Thai et al., 2014). We therefore create bootstrapped sample datasets by resampling individual tumor time series. Following Ref. (de Graft Acquah, 2012), we then repeat the model selection procedure on the two best performing models for each sample, and report the selection rate of each model (i.e. how often a model is ranked first according to the BIC):

| Samples: | Diffusive (no cov.) | Linear (no cov.) |
| --- | --- | --- |
| full dataset | 1 | 0 |
| 10 samples | 0.75 | 0.25 |
| 20 samples | 0.775 | 0.225 |

The diffusive model is therefore selected for more than 3/4 of the bootstrapped samples, which confirms its selection.

#### Subgroup analysis

The statistical model introduced above allows for the introduction of covariates, as in Eq. (11). We introduce the sex of the patient (M/F) as a categorical covariate for the parameter *r*. For each of the four growth models, SAEM converges to prior distributions that correspond to a lower median growth rates *r* for women than men (i.e. *r*_pop,F_ < *r*_pop,M_). With the diffusive model, we find *ξ*_M_ = 0.664(0.59) with *ξ*_F_ = 0, which corresponds to a factor 1.9 for the ratio of the median rates. We also find that the log-likelihood is improved compared to the no covariate case. However, the improvement is not large enough to compensate for the introduction of the extra model-parameter *ξ*_M_, and the BIC does not improve, as can be seen from the second column of Table 1. In ‘Supplementary Table 2’, we report the AIC values for the same experiments. Using this less stringent criterion for model selection would select the M/F covariate split for some of the growth models. To lift this ambiguity, we proceed to the Wald and LR tests

| Model: | Wald (p-value) | LR (p-value) |
| --- | --- | --- |
| Linear | × (0.089) | × (0.0949) |
| Diffusive | × (0.26) | × (0.2617) |

Using the threshold *p* = 0.05 on the p-values, both the Wald and the LR tests reject the covariate model. Thus using the sex as a covariate does not bring a statistically significant improvement for inferring the lesion growth rates.

Our model also permits to introduce multiclass categorical covariates, such as the lesion location categories presented in ‘Supplementary Table 1’. In ‘Supplementary Table 3’, we illustrate this and test for the significance of this covariate. We find an indication that focal lesions might grow slower in long bones (humerus, femur and tibia), but the limited amount of data does not permit to draw clear conclusions.

### 3.3 Modeling global tumor load

#### Tumor load effective model

Using SAEM, we estimate the population parameters of a mixed-effect model without covariate, as presented in Sec. 2.1, with the different growth models introduced above [Eqs. (6)-(8)]. We test the diffusive model, which was selected for the lesion growth in the previous section, and should therefore hold at short times, together with the IKS model, which should hold when the dissemination process dominates and different power-laws for the crossover regime, with *B* > 3/2.

The resulting BIC values are presented in the first column of Table 2. We find that the IKS model performs much better than the diffusive model (mean BIC separated by 40 confidence intervals), confirming that the dissemination process plays a great role in the tumor load evolution. However, power-law models with *B*≥3, lead to further improved values of the BIC (mean BIC improved by 8 confidence intervals for *B* = 6.5), indicating that we are in the crossover regime. In ‘Supplementary Table 4’, we systematically look for the most appropriate effective model, and compare BIC values for different values of *B*. This selects *B* = 6.5 as the best effective model for this dataset.

**Table 2:** Mean and standard deviation of BIC values for different tumor load growth models, in two mixed-effect models: without covariate (first column) and with the patient’s sex as a covariate for *R* (second column).

| Model: | no covariate | M/F cov. for $R$ |
| --- | --- | --- |
| Diffusive $B = 3/2$ | 1757.92 (0.13) | 1755.52 (0.12) |
| IKS (exponential) | 1745.42 (0.16) | 1742.09 (0.12) |
| Power-law $B = 3$ | 1744.15 (0.14) | 1741.40 (0.23) |
| Power-law $B = 6.5$ | 1741.30 (0.18) | 1738.19 (0.11) |

#### Subgroup analysis

We now use the sex of the patient as a categorical covariate for the growth rate *R*, as in Eq. (11). The introduction of this additional population parameter permits to improve the BIC for each parametric model (see second column of Table 2), indicating that the patient’s sex is relevant for modeling the evolution of the tumor load. We further proceed to the Wald and LR tests for the three best performing models

| Model: | Wald (p-value) | LR (p-value) |
| --- | --- | --- |
| IKS (exponential) | ✓ (0.008) | ✓ (0.0116) |
| Power-law $B = 3$ | ✓ (0.012) | ✓ (0.0160) |
| Power-law $B = 6.5$ | ✓ (0.0074) | ✓ (0.0131) |

The Wald and the LR test p-values are all smaller than 5%, such that the covariate model is selected in all cases. The sex is thus a relevant covariate for the tumor load growth rate *R*, and in each model, we find *R*_pop,F_ < *R*_pop,M_. For the power-law with *B* = 6.5, we report *R*_pop,F_ = 2.9(1.3) × 10^−3^ month^−1^ and *ξ*_M_ = 1.47(0.55), such that the median of the male population, 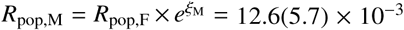 month^−1^, is four times as high. Those prior distributions are presented with the individual infered values in box ‘Tumor load modeling in SMM’. We also find *V*_0,pop_ = 1350(500) mm^3^, associated with the error parameter *b* = 0.281(0.027). The predictions from this model are shown in Fig. 5 for two patients, together with the initial lesion. Predictions for the whole cohort are displayed in ‘Supplementary Figure 2’. Further introducing the sex as a covariate for the initial volume *V*_0_ or for ω_*R*_, does not improve the BIC further, and gives negative Wald and LR tests, consistently over all growth models.

**Figure 5:**
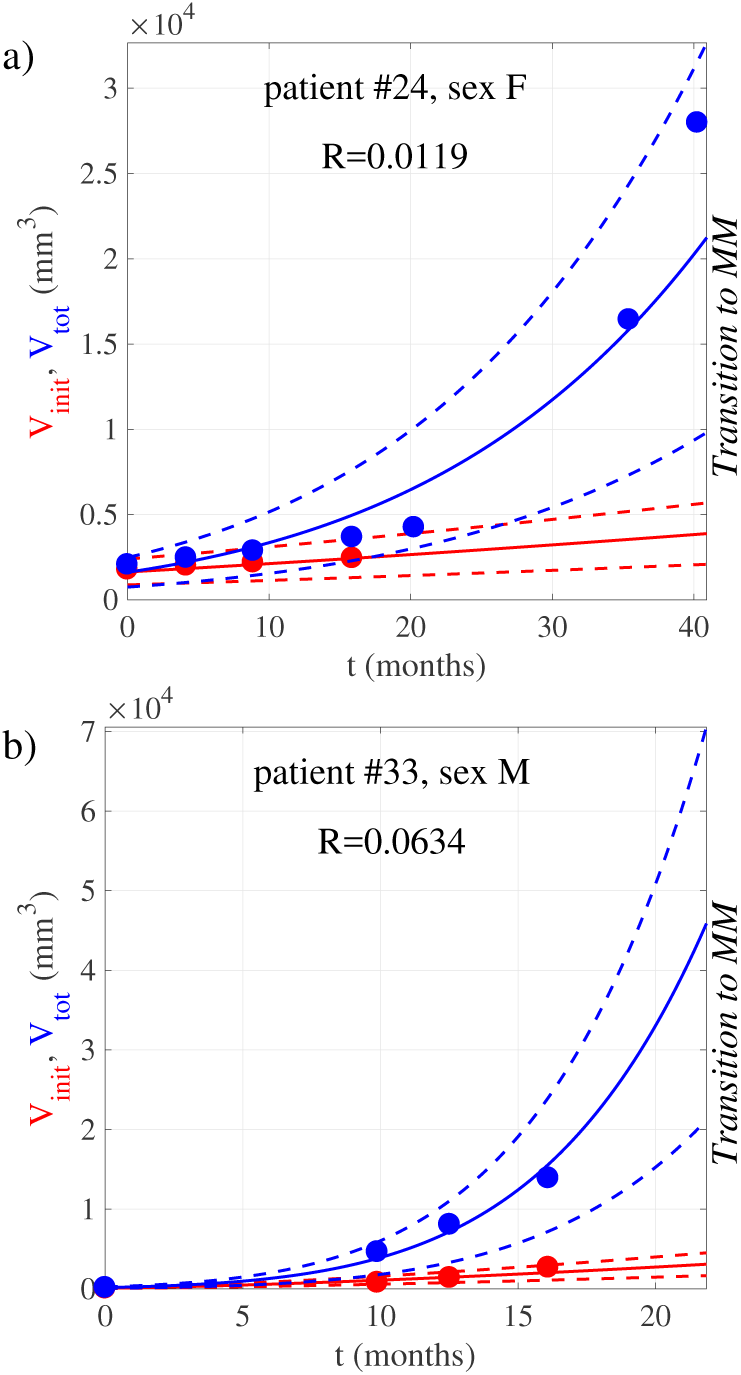
Tumor load *V*_tot_(*t*) (blue dots) and initial lesion *V*_init_(*t*) (red dots) time series with the predictions (solid lines) and the corresponding 95% error margin (dashed lines), for (a) a female and (b) a male patient, who both transition to MM. The predictions for the tumor load are based on the SAEM results using the power-law parametric model, Eq. (8) with *B* = 6.5, and a covariate model for the sex M/F.

###### Tumor load modeling in SMM

We find the effective model

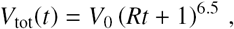

with the following median population values:

- *Growth rates for males and females* *R*_pop,F_ = 2.9(1.3) × 10^−3^ month^−1^ *R*_pop,M_ = 12.6(5.7) × 10^−3^ month^−1^
- *Initial volumes*: *V*_0,pop_ = 1350(500) mm^3^

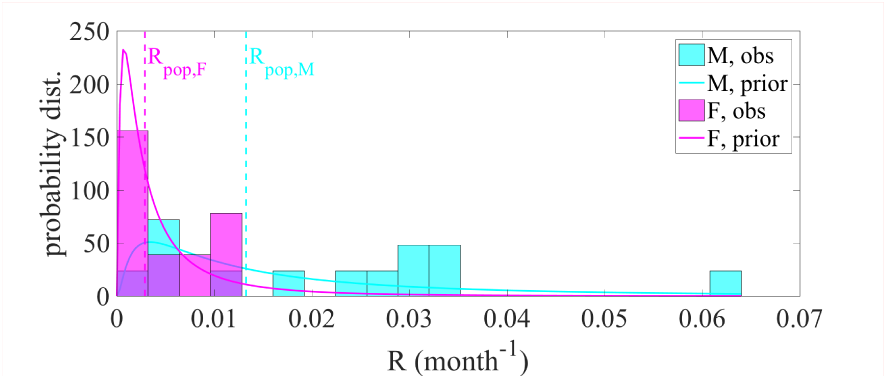

To assess the robustness of our model selection, we repeat the bootstrapping strategy from above and create bootstrapped samples, by resampling the tumor load time series. We repeat the model selection procedure, for each sample, and report the selection rate of each model, for the three best-candidates and using M/F as a covariate:

| Samples: | Power-law<br>$B = 3$ | Power-law<br>$B = 6.5$ | IKS<br>(exponential) |
| --- | --- | --- | --- |
| full dataset | 0 | 1 | 0 |
| 10 samples | 0.25 | 0.65 | 0.1 |
| 20 samples | 0.225 | 0.575 | 0.2 |

We see that the power-law model with *B* = 6.5 is selected in more than 50% of the cases, confirming our effective modeling approach for the crossover regime.

#### Dissemination

As a further check, we carried out a similar model selection analysis, applied to *N*_diss_(*t*), the number of disseminated lesions, which is expected to fully follow the IKS model Eq. (7), as it directly models the distribution of disseminated lesions. Using the total number of lesions, excluding the initial one, we compute the *N*_diss_(*t*) series and obtain a dataset of 13 patient series (8M and 5F) with 4.31 time points per series on average. We test in Table 3 the exponential model, and search for the best power-law model, in the model without covariate. We find a clear selection of the exponential model for *N*_diss_(*t*), thus nicely complementing our analysis of the crossover regime. In ‘Supplementary Table 5’, we further introduce the sex of the patient as a covariate. This analysis is, however, not conclusive: the model predicts a high ratio of the median of the rates in the two populations, but the Wald and LR tests do not permit to assert that those distributions are undistinguishable on this cohort.

**Table 3:**
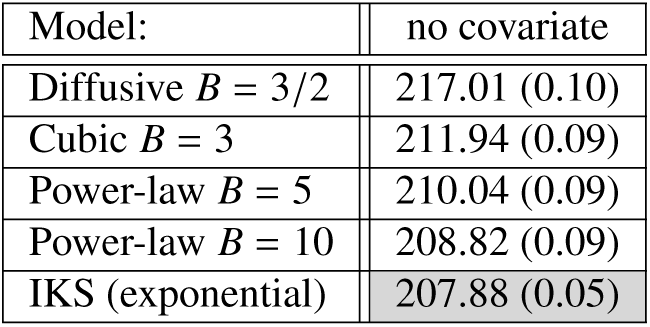
Mean and standard deviation of BIC values for the number of disseminated lesions, in a mixed-effect model with no covariate, (see ‘Supplementary Table 5’ for more values of *B*).

| Model: | no covariate |
| --- | --- |
| Diffusive $B = 3/2$ | 217.01 (0.10) |
| Cubic $B = 3$ | 211.94 (0.09) |
| Power-law $B = 5$ | 210.04 (0.09) |
| Power-law $B = 10$ | 208.82 (0.09) |
| IKS (exponential) | 207.88 (0.05) |

### 3.4 Novel model-based biomarkers

Based on this careful mathematical modeling of the disease evolution, that is agnostic of the survival chances of the patient, we propose to use patients’ parameters as model-aware biomarkers for clinical use. We show below that the tumor load growth rate *R* provides a relevant risk-stratification for MM. We then compare it with other non-model-based radiological biomarkers, and find indications that *R* is more relevant than the other criteria.

#### Transition to MM

As described in Sec. 3.1, we select the patients with at least two focal lesion measurements (26 patients), and include the information on progression to MM. For those patients, we compute the tumor load growth rate *R*, using the best performing model and priors from above i.e. power-law with *B* = 6.5 and M/F as a covariate, and the resulting prediction curves are shown together with the measured data in Fig 6. Using a threshold *R*_th_, we then use *R* to stratify patients into a low-risk (*R* < *R*_th_) and a high-risk (*R* ≥ *R*_th_) group to progress to MM. For each possible value of *R*_th_, we then compute the true positive rate for the detection of patient who progress to MM during the observation time, and the false positive rate, measuring the false alarms, and report them in the Receiver Operating Characteristic (ROC) curve (Zweig and Campbell, 1993), red line in Fig 7(a). Using the median of all observed growth ratesR as the stratification threshold, we obtain a true positive rate of 0.75 and a false positive rate of 0.1, as indicated by the red star. Considering the time of progression to MM, we show in Fig 7(b) the associated Kaplan-Meier curve (Kaplan and Meier, 1958). We compute the significance of the split with the log-rank test (Peto et al., 1977) and find a p-value of 0.00071, showing that the group compositions are statistically different. In ‘Supplementary Figure 3’, further searching for the best threshold in this population, we find that *R*_th_ = 7.7 × 10^−3^ month^−1^ gives a better p-value of 0.00003. We observe, however, that this precise threshold value might be overfitted to this dataset, and that all splits *R*_th_ ∈ [1.7 × 10^−3^, 4 × 10^−2^] month^−1^ consistently give p-values lower than 0.05. *R* is therefore a very relevant biomarker to predict the transition to MM.

**Figure 6:**
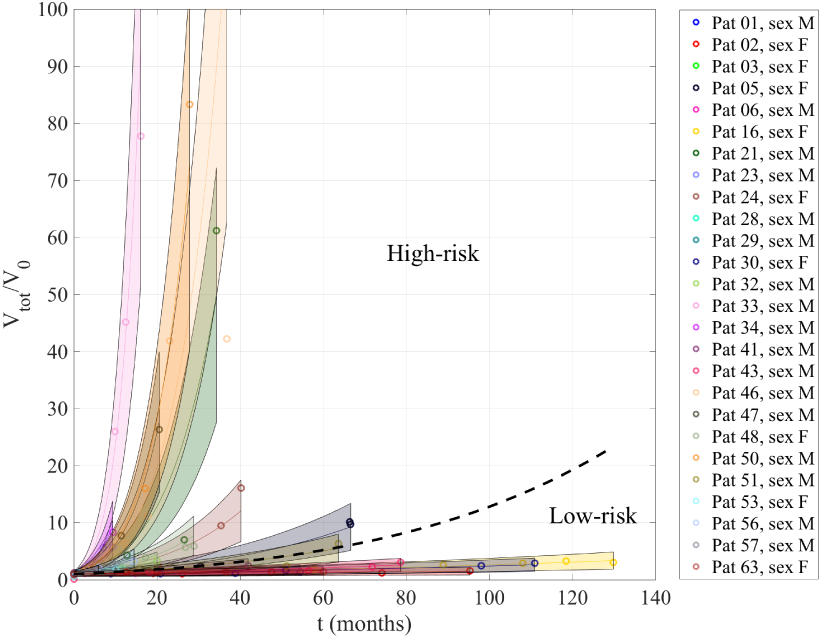
Rescaled tumor load series for patients with at least two time points, together with their fit using the best tumor load model and priors. The proposed risk groups are defined by the dashed separatrix.

**Figure 7:**
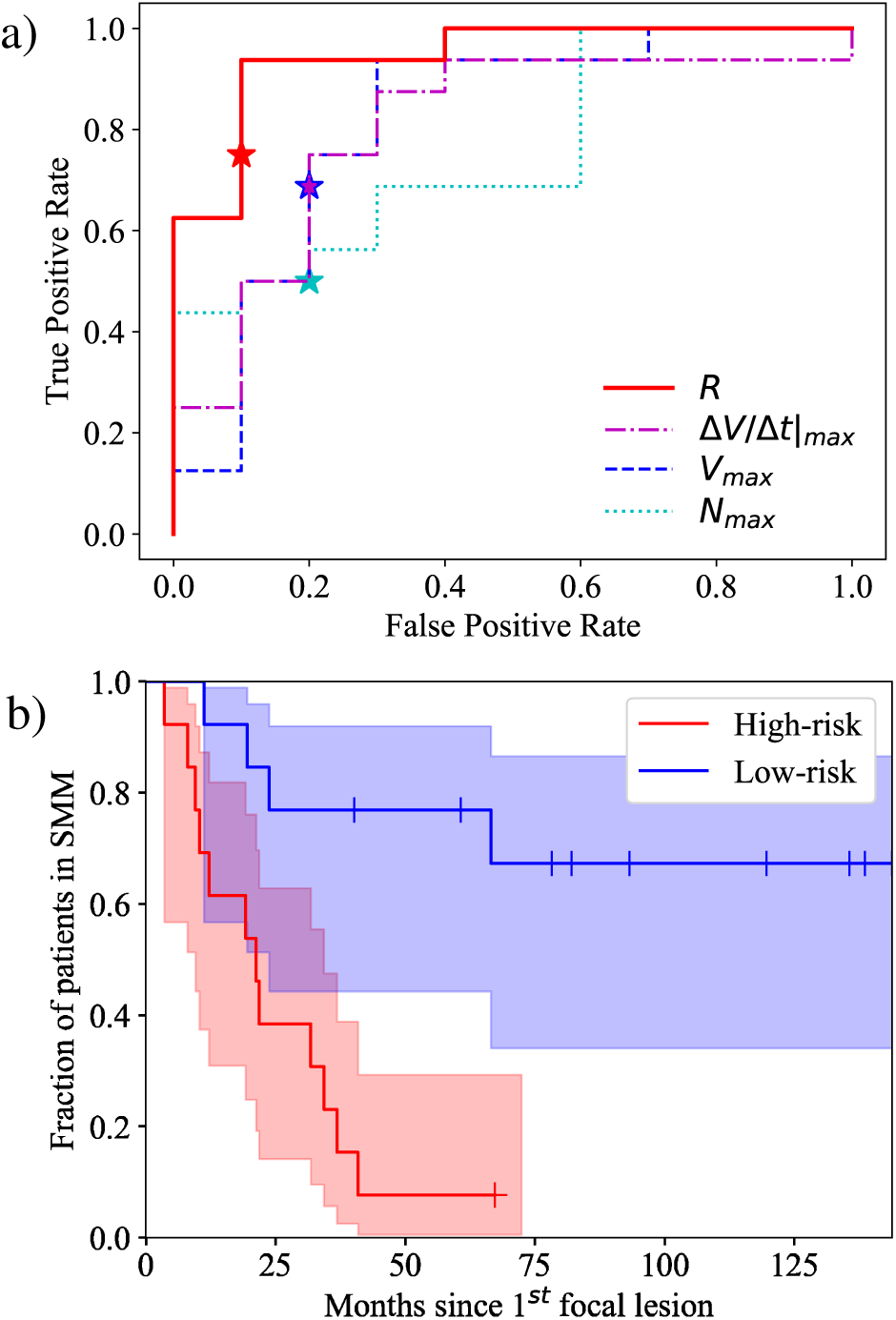
(a) ROC curve for different biomarkers, the red line corresponds to *R*. Stars indicate median splits, that are used in (a), (c) and Table 4. (b) Kaplan-Meier curve obtained for splitting this population into 2 groups using the median rate *R*_th_ = 1 × 10−2 month−1.

#### Comparison with other radiological criteria

We now compare this stratification with other MRI-based biomarkers. Previous studies (Hillengass et al., 2010; Merz et al., 2014; Brandelik et al., 2018; Wennmann et al., 2018) proposed radiological biomarkers for the risk stratification of SMM patients. Ref. (Brandelik et al., 2018) showed that volumetric measurements of tumors leads to a better assessment of the tumor load than diametric size, and Refs. (Merz et al., 2014; Wennmann et al., 2018) proposed to take into account the evolution between two measurements, by e.g. considering the rate of change of the tumor load (Wennmann et al., 2018). Here we therefore consider the following non-model-based biomarkers, which, for a fair comparison with the proposed biomarker *R*, are also volumetric and retrospectively based on all available measurements:

**Table 4:** Area Under the Curve values of the ROC curves from Fig. 7(a), Kaplan-Meier log-rank test p-values and relative risk for progression to MM, associated to the stratification into low- and high-risk groups when using the median observed value of different radiological criteria. Details on those tests for evaluating the discriminative power of biomarkers are given in ‘Supplementary Method 2’.

|  | Criteria | Median value | Area Under the Curve | p-value | relative risk |
| --- | --- | --- | --- | --- | --- |
| I | $N_{\max}$ | 4 lesions | 0.75 | 0.2038 | 2.39 |
| II | $V_{\max}$ | 8816 mm<sup>3</sup> | 0.81 | 0.0116 | 5.28 |
| III | $\Delta V/\Delta t\vert _{\max}$ | $5.2 \times 10^{-1}$ mm<sup>3</sup> month<sup>-1</sup> | 0.80 | 0.0130 | 5.13 |
| IV | $R$ | $1.0 \times 10^{-2}$ month<sup>-1</sup> | 0.94 | 0.0007 | 9.58 |

I. - the largest observed focal lesions number, 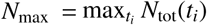,
II. - the largest observed tumor load, 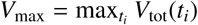,
III. - the largest rate of change of the tumor load be-tween two consecutive measurements, 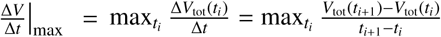.

Figure 7(a) displays the ROC curves for those criteria, and we see that they provide a fair stratification as well, although not as good as the one provided by *R*. In Table 4, we report the corresponding Area Under the Curve values, which reaches 0.94 for *R*. We also report the log-rank test p-values associated to the Kaplan-Meier survival curves obtained using each of those criteria and taking the median observed value as stratification threshold, together with the associated relative risk (Stare and Maucort-Boulch, 2016). Criteria ii and iii are both providing a statistically relevant classification, with comparable p-values ∼ 0.01 and comparable relative risks ∼ 5.1 − 5.3. In row iv, we see that the growth rates *R* – i.e. the model-based approach – provide the most relevant split, with a significantly lower p-value = 0.0007 and the best risk-stratification relative risk = 9.6, compared to other radiological criteria.

## 4. Discussion

### Diffusive growth of local lesions

In Sec. 3.2, we analyzed disease progression at the local scale. We found that the growth of focal lesions in MM is best modeled by a diffusive growth, thereby confirming a basic hypothesis of Ayati *et al.* (Ayati et al., 2010) and is in line with phenomenological observations, in particular that lesions sometimes become too diffuse to be properly volumetrized. This could be further analyzed by the study of the evolution of lesion borders, possibly in multiple imaging modalities (Konukoglu et al., 2010; Lipkova *et al*., 2018) With respect to other tested models, we found that cubic growth, which would correspond to a solid tumor with a rate of growth proportional to the tumor surface, performed better than exponential growth. This does not come as a surprise, as the latter would correspond to a volumetric growth, implying that the newly generated mass is ‘pushing’ its surrounding, which may be rather unlikely to happen in a bone environment.

### Effective model for the crossover to global dissemination

In Sec. 3.3, we considered the global propagation of the disease on the patient’s scale, through the analysis of the tumor load. We showed that the diffusive growth parametric model does not translate to the tumor load as such. Here, a model taking the dissemination process into account, like the IKS model using an exponential term (Iwata et al., 2000), is likely more relevant in our SMM cohort, where most of the monitored patients show a progress in the number of lesions observed. Still, the tumor load model that aligns best with our observations is a powerlaw with *B* = 6.5. As summarized in box ‘Descriptive tumor load model’, we interpret this as a crossover from an initial diffusive regime (power-law with power 3/2) to an exponential dissemination regime. This is best effectively modeled by a power-law model with a higher power *B* > 3/2. Further tests on the number of disseminated lesions confirm that the basic assumptions of the IKS model (Iwata et al., 2000), i.e. exponential dissemination, apply in this cohort.

In the original paper, the IKS model was validated on one single patient with a metastatic hepatocellular carcinoma (Iwata et al., 2000) and has been further tested on one other patient with liver cancer and one with lung cancer in Ref. (Mehrara et al., 2013). A few population studies have been carried out on mice populations with orthotoptic cell implantation (Hartung et al., 2014; Baratchart et al., 2015)(Benzekry et al., 2016), the observed dissemination dynamics also showing an overall good agreement with the IKS model, although Ref. (Baratchart et al., 2015) incorporated interaction between lesions growing in close vicinity to match the experimental conditions. The IKS model has also been used in breast cancer to fit cross-sectional data on the risk of metastatic evolution (Barbolosi et al., 2011) using ad hoc parameters, and Ref. (Benzekry et al., 2016) used it to predict metastatic relapse. Another study of brain metastasis in non-small cell lung cancer (Bilous et al., 2018) was concomitant to ours. We are, however, not aware of any previous longitudinal human population study of the IKS model in the context of MM, such that our study, with 21 patients in the crossover regime and 13 series for the number of disseminated lesions, is unprecedented.

### Role of the sex and other covariates

Our hierarchical statistical model is evaluated on the whole population jointly, and permits to systematically test for the impact of covariates on model parameters. We found that the patient’s sex is a relevant covariate to predict the tumor load growth rate, with a median rate four times as high in the male population as in the female one. The role of the patient’s sex in the incidence of MM is known (Raab et al., 2009; Roellig et al., 2015; Kumar et al., 2017), but no indication of its role in the disease evolution in the presence of focal lesions has been previously reported. However, it has been shown that activated estrogene receptors inhibit cell survival pathways and support cell apoptosis in MM (Sola and Renoir, 2007), which provides one possible explanation. This effect deserves to be further investigated in a larger cohort. The sex covariate alone is, however, a less relevant covariate for the local lesion growth modeling, indicating that other hidden covariates might play a role and could be added in the model as well. The location of focal lesions could be a candidate, and in ‘Supplementary Table 3’ we have found indication that long bones tend to have lower growth rates than other bones. Our sample size is, however, too small to carry out multi-covariate tests with enough statistical strength. The role of the sex covariate in tumor load modeling in our cohort could also be explained by a difference in the dissemination rate, but more statistical strength would be needed to conclude here as well.

### Growth rates as model-based biomarkers

In Sec. 3.4, we identified the tumor load growth rate as viable image biomarker that is integrating information along the full observational sequence. We have shown that it provides a pertinent risk-stratification of SMM patients to develop end-organ damage and therefore transition to MM. This biomarker provides a better risk-stratification than other MRI-based biomarkers that have been suggested in the literature, even when few observations are present, gaining strength from the population priors. The biomarker gets refined over time when the number of observations increases, as it takes all available measurements into account, thereby confirming the unprecedented potential of model-based biomarkers for better and more personalized treatment decisions. This contrasts with current biomarkers, which consider the most recent examination only (International Myeloma Working Group, 2003; Rajkumar et al., 2014; Rajkumar, 2016). The model used for the biomarkers was derived on part of the population on which the biomarker is tested. This however does not lead to overfitting, as the growth model and the model selection process do not know about survival and progression to MM. We do acknowledge the limited size of the patient’s cohort and the retrospective nature of the study, and for full clinical relevance, the proposed model-based radiological biomarkers should be combined with the remainder of MM biomarkers in future studies.

## 5. Conclusion

In this paper, we propose a descriptive functional-statistical framework to carry out multi-scale modeling of cancer evolution, from single lesion growth to global dissemination. Applying it to MM, we tested different mathematical models, which permits to confirm basic assumptions on disease progression (Ayati et al., 2010; Iwata et al., 2000). We learned population priors, as well as tested the influence of various covariates, on clinical data. Our study establishes a new benchmark for the study of metastatic and disseminated diseases in general, and for understanding the progression of MM in particular. We also propose to use the inferred model parameters as biomarkers, and showed the relevance of the growth rate to predict the transition to overt MM in our dataset. We were able to show that biomarkers based on biologically-grounded, but tractable models could be more significant than phenomenological ones, which offers new and unprecedented directions.

Model-based biomarkers could be used in the clinical routine, as inference of the individual parameters is very fast. Indeed, for our study, it takes 0.15s per patient on average with the Monolix software (Monolix). This computation could therefore be implemented after computeraided segmentation of focal lesions in MRI, for example. The use of whole-body imaging and whole-body lesion analysis is hindered by a lack of means to postprocess and analyze these added information. Our model now offers such means fostering the impact of high capacity data analytics in clinical decision making.

Future work should integrate other image-based and non-image-based, static and dynamical features in a radiomics approach. Our model could be extended and refined by considering the interaction between lesions in the growth process, such as done in (Baratchart et al., 2015; Benzekry et al., 2017) One could in particular investigate related models for other observed parameters for MM, such as laboratory parameters that are known to correlate with the progression of the disease (Mai et al., 2015; Wennmann et al., 2018), genetic markers or cell-surface proteins measured with flow-cytometry (Flores-Montero et al., 2017). Next-generation sequencing, which is becoming available and permits to distinguish different clonal phenotypes of plasma cells (Takamatsu, 2017) would enable to analyse separately the role of different mutations in disease progression.

## Acknowledgments

The authors would like to thank the Deutsche Forschungsgemeinschaft (DFG; WE 2709/3-1 and ME 3511/3-1) and the Austrian Science Fund (FWF; I2714-B31) for research funding. U.K. acknowledges founding from the Deutsche Forschungsgemeinschaft (grants SFB 824 and SFB 1335) and from the Deutsche Krebshilfe (grants 111305 and 111944), and is further supported by Stiftung Charité.

## Supplementary Material

### Supplementary Method 1. Model comparison

We present here the different metrics that we use to qualitatively assess the adequacy of a model, and perform model selection.

### Akaike and Bayesian Information Criteria

The Akaike Information Criterion (AIC) is defined as (Akaike, 1973)

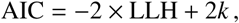

where *k* is the total number of parameters, i.e. the length of the vector which gathers all population and error parameters. The Bayesian Information Criterion (BIC) in turn reads (Schwarz, 1978)

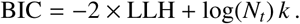

Both criteria are meant to be minimized, hereby tending to maximize the log-likelihood, i.e. the goodness of the fit. They both contain a penalty term for models which have a greater number of free parameters, in order to select models that are more efficient in modeling the data. We see that the BIC penalizes model complexity more heavily as soon as *N*_*t*_ > 7. Comparison of the performance of those criteria shows that AIC performs better for small number of time series *N*_*t*_, whereas the BIC is more consistent for larger datasets (Markon and Krueger, 2004; de Graft Acquah, 2012).

### Wald and Likelihood ratio tests

In the case where one model is a particular case of a more complex one, there exists further statistical tests for model comparison. This applies in particular to testing the relevance of the use of a covariate in a model, as a model with covariates embeds the simpler case without covariate [e.g. take *ξ*_*k*_ = 0 for *k* ∈ [1, *K* − 1] in Eq. (11)]. The Wald test (Wald, 1943) aims at testing if the inferred distributions of the extra-parameters [{*ξ*_*k*_}_*k*=1..*K*−1_ in Eq. (10)] of the more complex model are statistically distinguishable from the value they take in the simpler model, in the so-called null hypothesis. Those distributions are approximated by normal ones, whose variance is estimated via the Fisher information matrix, and the corresponding p-value computed to assess the plausibility of the null hypothesis. The likelihood ratio (LR) considers the ratio of the likelihood of the two models (Neyman and Pearson, 1933). The distribution of its logarithm is then compared to a chi-square distribution, which is expected in the null hypothesis (Wilks, 1938), via the computation of the p-value. The Wald approach is simpler, and asymptotically equivalent to the LR test. It is, however, less reliable in some cases, in particular as it cannot capture any structure or asymmetry of the likelihood function (Meeker and Escobar, 1995; Pawitan, 2000).

### Supplementary Method. Biomarker evaluation

We introduce here different methods to assess the relevance of a biomarker in risk-stratification. We are concerned in this study with binary clinical outcome, i.e. whether the patient progresses to MM or not.

### Receiver Operating Characteristic (ROC) curve

The ROC curve is often used in the clinical medicine context (Zweig and Campbell, 1993). In this case, each test time series is associated with a value of the biomarker, as well as a positive or a negative label, labeling if the event (progession to MM) occurred or not over the observation period. For a split in a high-risk and a low-risk group, we compute the True Positive Rate (TPR), measuring the probability that a positive example was classified in the high-risk group, and the False Positive Rate (FPR), measuring the rate of negative events classified in the high-risk group:

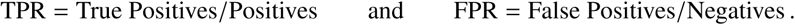

The ROC curve consists in plotting the TPR against the FPR for various thresholds of the biomarker. A ROC curve is considered discriminative if it is above the TPR=FPR line, that would correspond to random assignments to the risk-groups. The discriminative power of a biomarker can be assessed by the area under the ROC curve, which should be above 0.5 and as close to 1 as possible; a value of 1 corresponding to a perfect discriminator.

### Kaplan-Meier survival plots

Risk prediction in time can be made using Kaplan-Meier ‘survival’ plots (Kaplan and Meier, 1958), which take the timing of the event into account. This plot estimates the survival function for each group, taking into account censored data (i.e. when the observation ends, without occurrence of the event). The log-rank test measures if the survival distributions of the two groups are statistically distinguishable (Peto et al., 1977). To quantify the difference in survival between the two groups, we compute the Relative Risk (Stare and Maucort-Boulch, 2016):

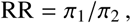

where *π*_1,2_ are the probabilities of event occurrence in each group, i.e. the total follow-up time, divided by the number of events.

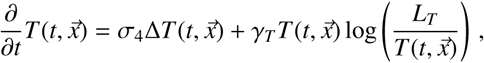

### Supplementary Method 3. Derivation of the diffusive growth model

Reference (Ayati et al., 2010) assumes a diffusion equation with a Gompertz-like saturation term for the tumor density, in one-dimension (see Eq. (33) in Ref. (Ayati et al., 2010)). We extend it to three dimensions in an isotropic manner:

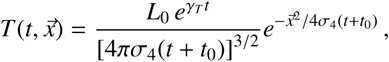

where 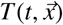 is the local tumor density, *γ*_*T*_ > 0 is the Gompertz growth constant, and *L*_*T*_ the maximum tumor density, and *σ*_4_ is the diffusion coefficient of the lesion. In the lesion area (i.e. where 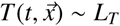, the growth is essentially diffusive, with a reaction term that can be simplified to:

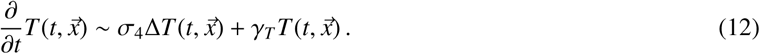

Solving Eq. (12) with an initial isotropic lesion of size *d*_0_, using 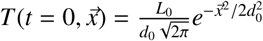, we find the evolution of the tumor density

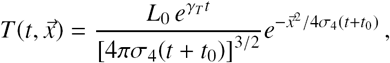

with 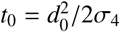. Computing the linear size of the lesion using the root-mean-square distance from the origin, gives

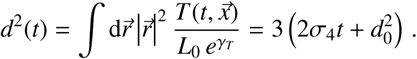

The time evolution of the associated volume is therefore 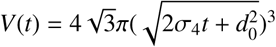. After reparametrization, we find

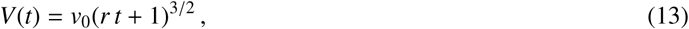

which corresponds to 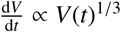.

### Supplementary Method 4. Solutions of the IKS model

The IKS equation models the size distribution of disseminated lesions in time ρ(*v, t*) (Iwata et al., 2000). This quantity follows a transport equation:

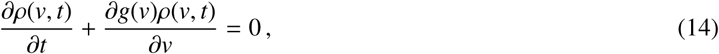

where *g*(*v*) is the growth rate of individual lesions 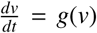. Considering the standard boundary conditions, also presented in Ref. (Iwata et al., 2000; Struckmeier, 2003; Evys et al., 2009), one can solve those equations.

Full analytical solution of Eq. (14) is obtained in the case of exponential growth for the individual lesions (i.e. *g*(*v*) = *rv*) which is for example presented in Ref. (Struckmeier, 2003):

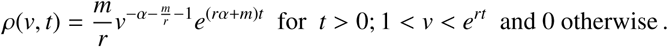

In the cases of the power-law lesion growth models, we use *g*(*v*) = *av*^1−*γ*^, where *γ* = 1 holds for a linear growth, 1/3 for a cubic growth and 2/3 for the diffusive growth. In equation (16), one then finds 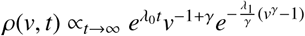, with *λ*_*0*_ = *aλ*_*1*_ and where *λ*_*1*_ is the maximum real solution of the eigenvalue equation 1 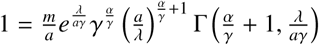 (Iwata et al., 2000).

Therefore, for the different lesion growth models described in Sec. Lesion growth models, the distribution ρ(*v, t*) is asymptotically separable (Evys et al., 2009), and has an exponential time-dependence

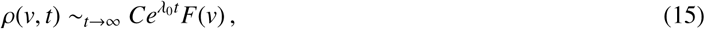

where λ_0_ and *F* depend non-trivially on the growth and the dissemination models. This will translates into an exponential asymptotic behaviour for both the number of disseminated lesion *N* (*t*) *dv* ρ(*v, t*) and the disseminated burden *V*_diss_(*t*) = ∫ *dv v*ρ(*v, t*):

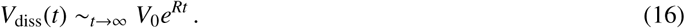

**Supplementary Table 1.** Lesion location categories in the individual lesions dataset. This table presents the distribution of the locations of lesions in the individual lesions dataset, amongst time series and patients.

|  | Category | Number of lesions | Number of patients |
| --- | --- | --- | --- |
| 1 | Skull | - | - |
| 2 | Cervical vertebrae | 2 | 1 |
| 3 | Thoracic vertebrae | 10 | 4 |
| 4 | Lumbar vertebrae | 6 | 4 |
| 5 | Sacrum & Coccyx | 5 | 5 |
| 6 | Shoulders | 2 | 2 |
| 7 | Humerus | - | - |
| 8 | Pelvis | 7 | 6 |
| 9 | Femur | 12 | 8 |
| 10 | Tibia & Foot | 2 | 1 |
| 11 | extra-osseous | 1 | 1 |
| 12 | Ribs | - | - |
| 13 | Sternum | 2 | 2 |

**Supplementary Table 2.**
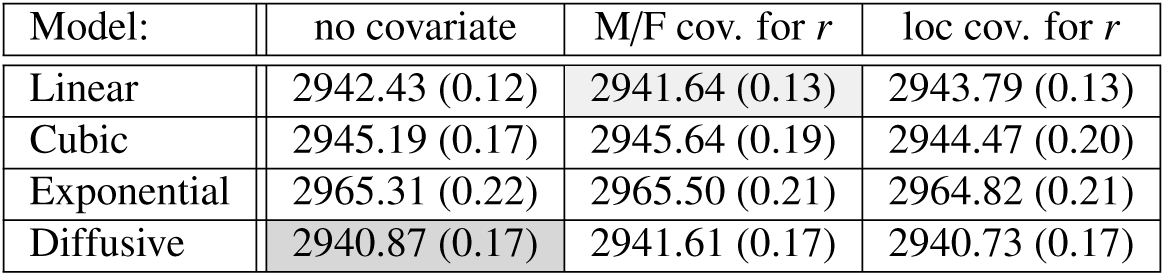
AIC values for lesion growth model selection. We present the AIC values for the different lesion growth models, with three mixed-effects models: without covariate (first column), with the patient’s sex as a covariate for *r* (second column), and with the lesion location as a covariate for *r* (third column), as detailed in Supplementary Table 3.

| Model: | no covariate | M/F cov. for $r$ | loc cov. for $r$ |
| --- | --- | --- | --- |
| Linear | 2942.43 (0.12) | 2941.64 (0.13) | 2943.79 (0.13) |
| Cubic | 2945.19 (0.17) | 2945.64 (0.19) | 2944.47 (0.20) |
| Exponential | 2965.31 (0.22) | 2965.50 (0.21) | 2964.82 (0.21) |
| Diffusive | 2940.87 (0.17) | 2941.61 (0.17) | 2940.73 (0.17) |

*Supplementary Table 3. Lesion location covariate testing.* Using the lesion location categories (see Supplementary Table 1), we test for the lesion location as a multiclass covariate in the lesion growth model. However, as some categories are not, or almost not, represented in our dataset, we group different locations together. We obtain a low log-likelihood when grouping the lesions of the spine in one group (categories 2, 3, 4 and 5 in Supplementary Table 1, denominated ‘spine’ group in the following), those in long bones in a second (categories 7, 9 and 10, denominated ‘long’ group) and the remaining categories together (categories 1, 6, 8, 11, 12 and 13, denominated ‘0’ group). Long bones are known to have a different trabecular structure than other bones. Introducing those groups as a 3-fold covariate for *r* in the diffusive model, we find that SAEM converges to lower growth rates for the long bones group, i.e. *r*_pop,long_ < *r*_pop,0_, with *ξ*_leg_ = − 0.749(0.68), and to growth rates for the spine group that are not distinguishable from the other bones, with *ξ*_spine_ = 0.0607(0.64).

We therefore further analyze the binary split ‘long’ vs {‘spine’ and other bones}. The log-likelihoods are comparable to that of the ternary split and the BIC values are reported below:

| Model: | loc cov. for $r$ |
| --- | --- |
| Linear | 2955.14 (0.13) |
| Cubic | 2955.82 (0.20) |
| Exponential | 2976.17 (0.21) |
| Diffusive | 2952.09 (0.17) |

We find that the BIC increases compared to the no covariate case (first column of Table 1). The AIC criterion, however, would select this covariate split in some cases, and does not permit to decide within the error bars for the diffusive model (see third column of Supplementary Table 2). To confirm, we again proceed to the Wald and LR tests:

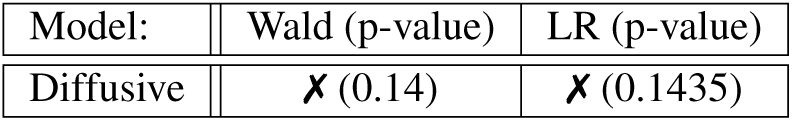

Both the Wald and the LR tests reject the covariate model. Thus using the ‘long’ bone covariate does not bring a significant improvement for inferring the local lesion growth rates on this dataset. However, we found an indication that tumors might grow slower in those bones. We also observe that the p-value of those tests tends to decrease with the dataset size, indicating that a larger cohort could permit to draw significant conclusions.

**Supplementary Table 4.** Effective model for the tumor load. We perform model selection for the tumor load time series, comparing power-law models [Eq. (8)] with different values of *B*. We estimate, with SAEM, the population parameters of a mixed-effect model without covariate and with the sex of the patient (M/F) as a categorical covariate for the growth rate *R*, as in Eq. (11). The BIC values are gathered in the following table:

| $B$ | no covariate | M/F cov. for $R$ |
| --- | --- | --- |
| 0.5 | 1792.83 (0.12) | 1789.86 (0.11) |
| 1 | 1772.12 (0.14) | 1769.91 (0.11) |
| 1.5 | 1757.91 (0.24) | 1755.52 (0.12) |
| 3 | 1744.19 (0.14) | 1741.40 (0.23) |
| 5 | 1741.38 (0.15) | 1738.58 (0.22) |
| 5.5 | 1741.27 (0.19) | 1738.42 (0.21) |
| 6 | 1741.35 (0.16) | 1738.30 (0.19) |
| 6.5 | 1741.22 (0.15) | 1738.19 (0.11) |
| 7 | 1741.24 (0.16) | 1738.23 (0.23) |
| 7.5 | 1741.32 (0.14) | 1738.39 (0.15) |
| 8 | 1741.45 (0.14) | 1738.50 (0.16) |
| 10 | 1741.89 (0.15) | 1738.78 (0.23) |
| 12 | 1742.18 (0.18) | 1739.02 (0.13) |
| 20 | 1743.31 (0.18) | 1739.95 (0.21) |

This permits to select the model with *B* ∼6.5 as the best effective model.

Considering *B* as a patient-dependent parameter in Eq. (8), we also run SAEM with a normal prior on *B*_*j*_, i.e. *B*_*j*_ = *B*_pop_ + η _*j*_ with 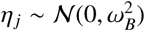, and we fix ω_*B*_ = 2. We use the sex of the patient as covariate for *R*, as selected by the previous experiment. The value selected by the algorithm for the parameters of the prior is *B*_pop_ ∼ 6.39(1.3), corresponding to a BIC of 1741.58(0.15). This population value is fully consistent with the best value of *B* ∼ 6.5 selected above. Introducing a patient-dependent width to the prior can be justified by the fact that we are analyzing an effective transitional behaviour, and different patient series might be at a different stages of the transition. However, this additional degree of freedom does not permit to reduce the BIC values.

**Supplementary Table 5.**
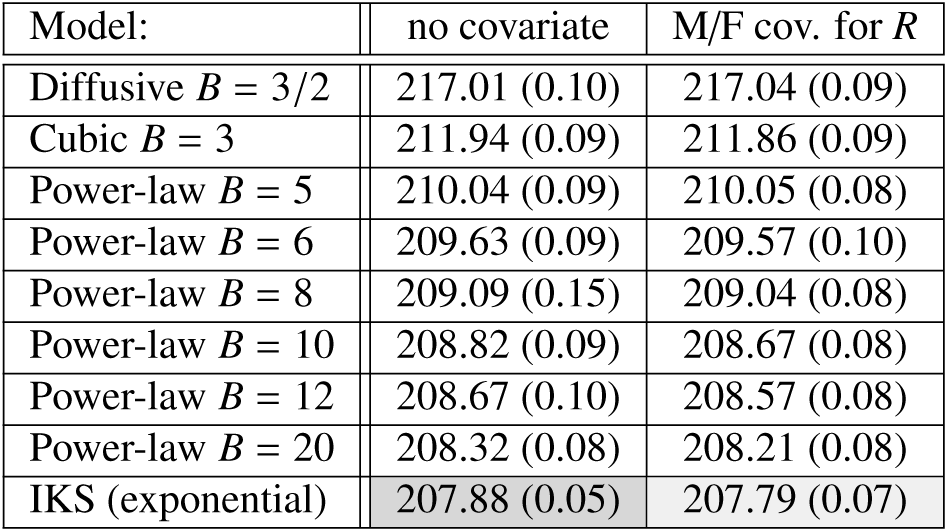
Model selection on the number of disseminated lesions. We proceed to an analysis similar to Sec. Modeling global tumor load, on *N*_diss_, the number of disseminated lesions, which is expected to fully follow the IKS model, Eq. (8). Computing the *N*_diss_(*t*) series (using total number of lesions, excluding the initial lesion), we obtain a dataset of 13 patient series (8M and 5F) with 4.31 time points per series on average. As in Supplementary Table 4, we test the exponential model, and we search for the best power-law model, using both no covariate, and the sex as a covariate for *R*. We compare the BIC values obtained with SAEM, on those experiments:

This clearly selects the exponential model over the power-law ones, confirming that the dissemination process follows the IKS model. The BIC is (very) slightly improved in the M/F covariate model for *R*, but both values are compatible within the error bars. In the covariate model, we obtain *ξ*_M_ = 1.98(1.3), which corresponds to a high ratio 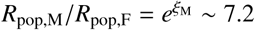 for the population median, but is associated to a very high variance. We therefore proceed to the Wald and LR tests, which both rejects the covariate model on this cohort, considering a 5% threshold.

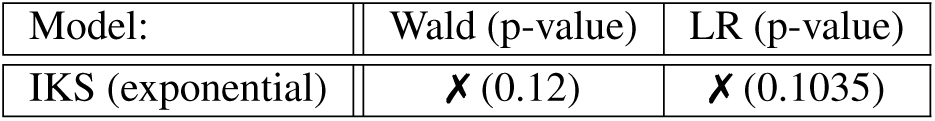

**Supplementary Figure 1.**
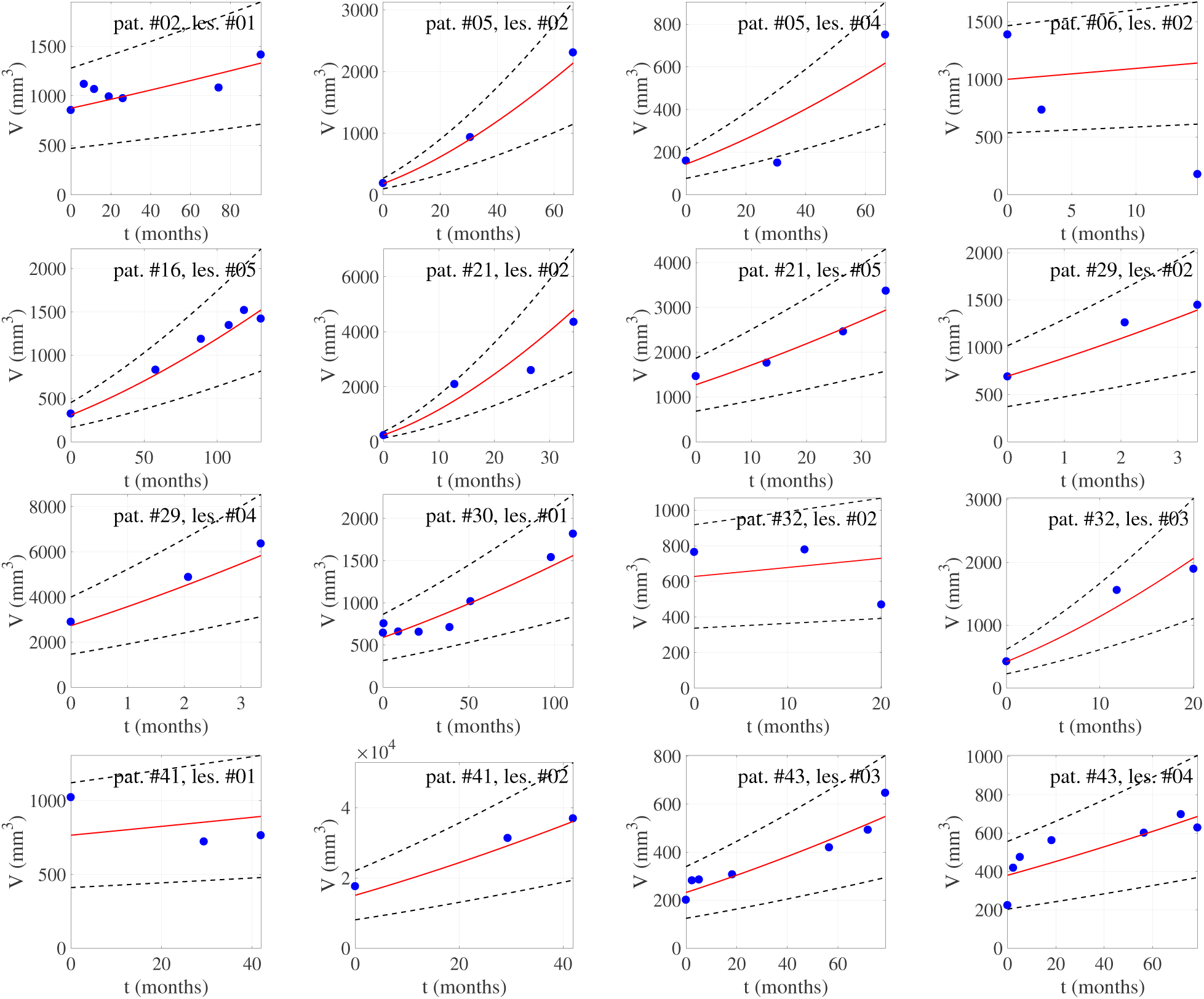
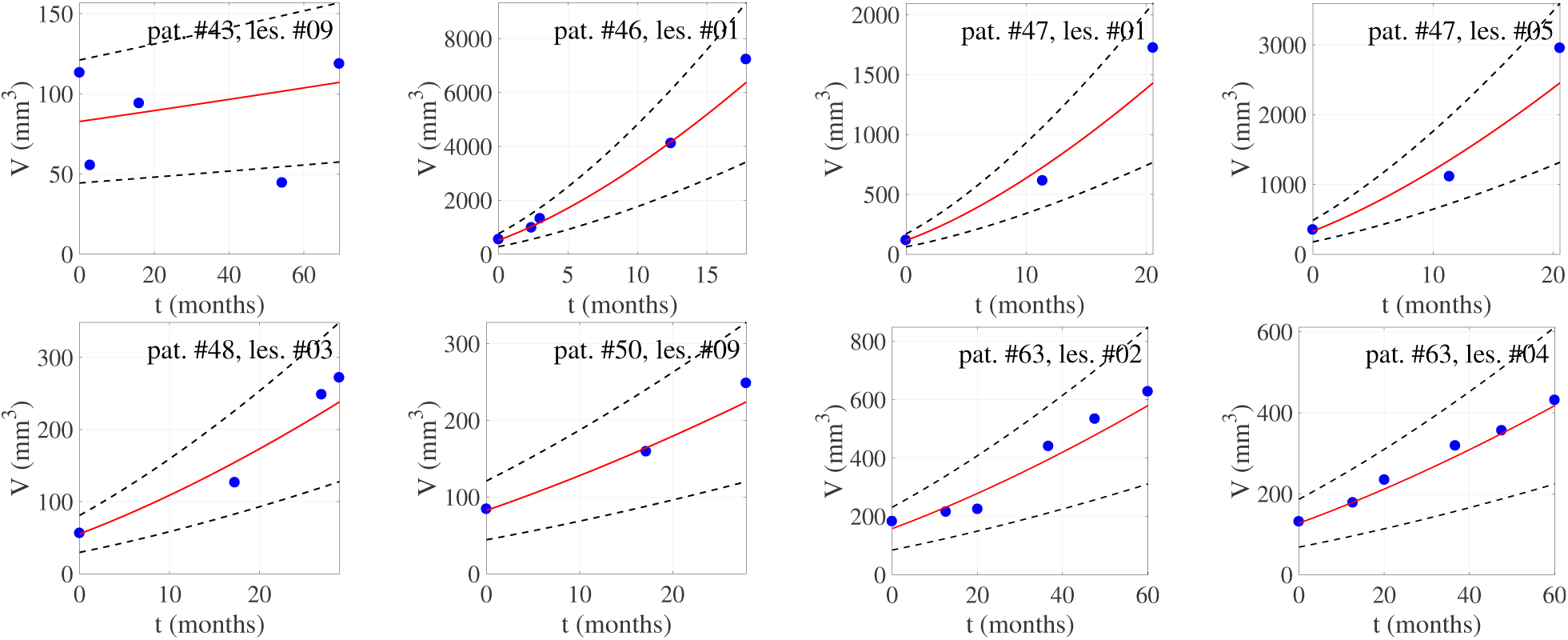
Focal lesion time series. We show here selected focal lesion growth series *V*(*t*) (blue dots) with individual predictions (red lines) and the corresponding 95% error margin (black dashed lines). The predictions are based on the SAEM results using the diffusive parametric model, Eq. (4).

**Supplementary Figure 2.**
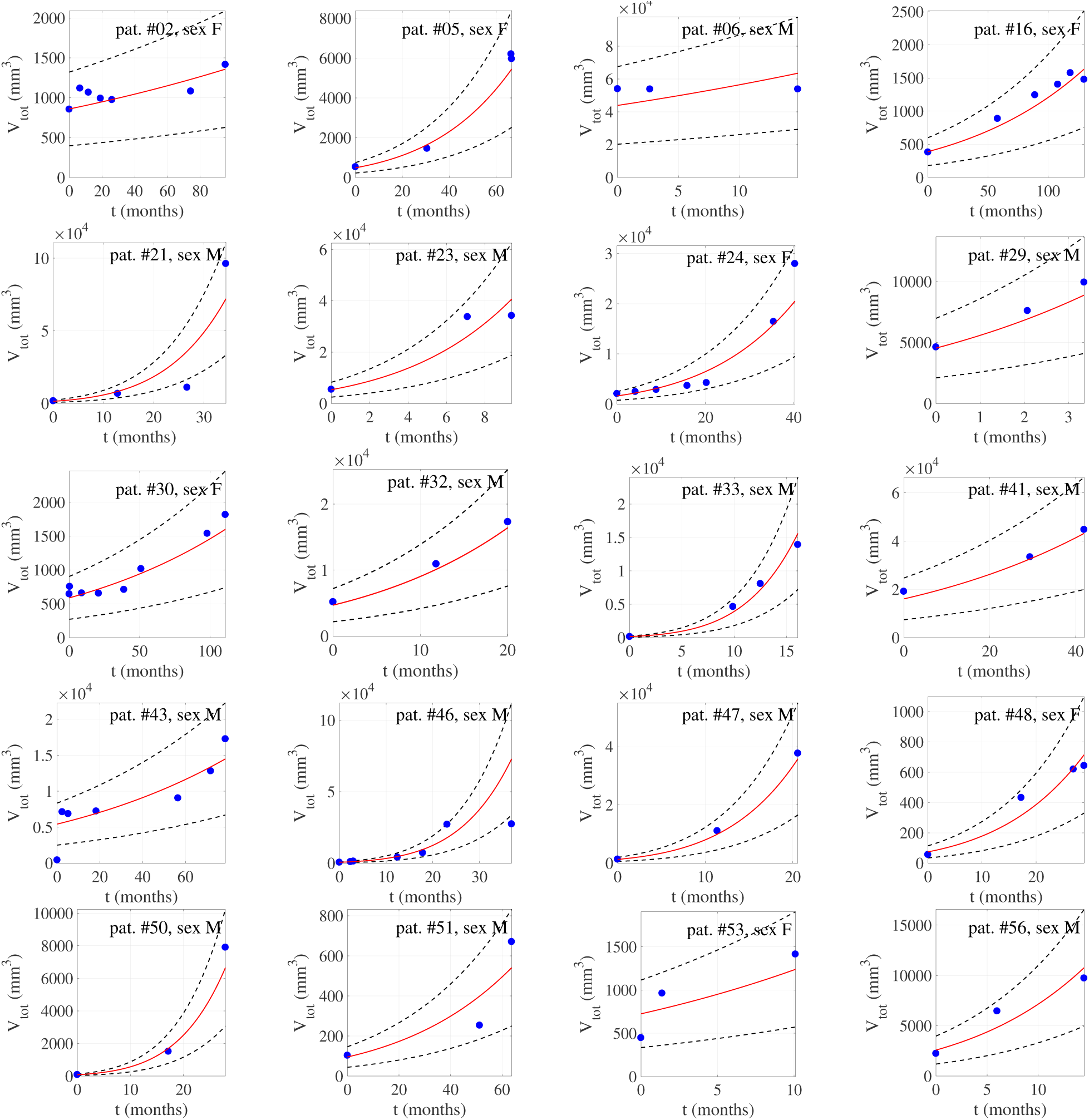
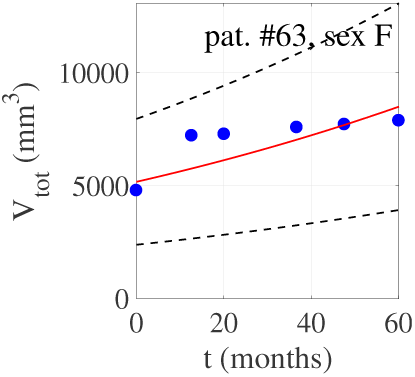
Tumor load time series. We show below the tumor load time series *V*_tot_(*t*) (blue dots) with individual predictions (red lines) and the corresponding 95% error margin (black dashed lines). The predictions are based on the SAEM results using the power-law parametric model, Eq. (8), with *B* = 6.5, and a covariate model for the sex M/F.

**Supplementary Figure 3.**
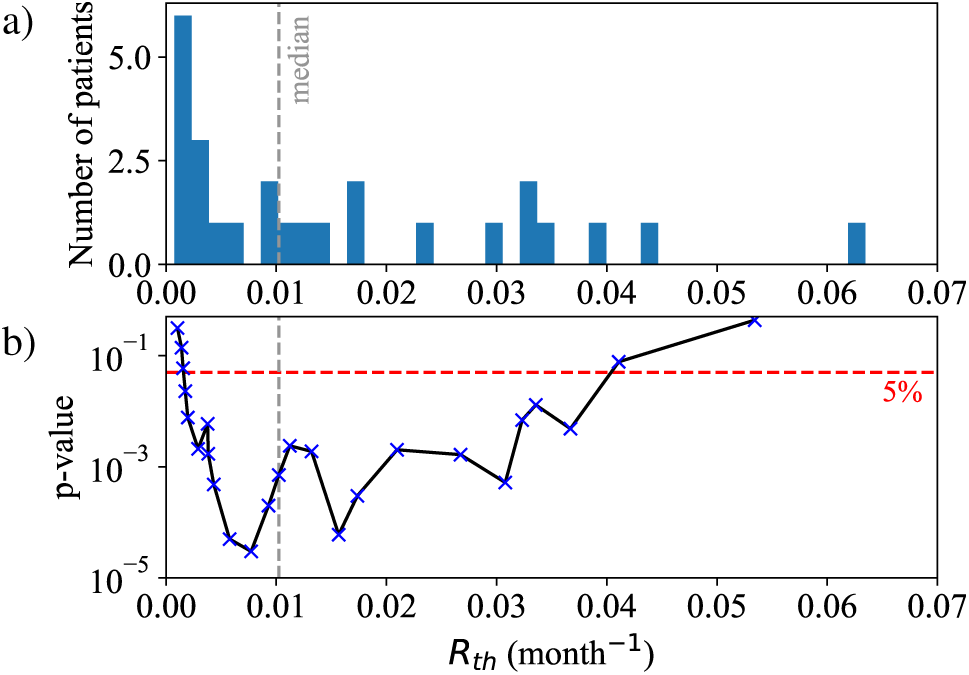
Varying the threshold R_*th*_. The figure below shows (a) the histogram of the tumor load growth rates *R*, used as a biomarker, (b) the Kaplan-Meier p-values obtained for different threshold values *R*_th_.

